# A human-in-the-loop approach for faster cell tracking

**DOI:** 10.64898/2026.10.06.756962

**Authors:** Mihaela Mihaylova, Ankur Ankan, Shabaz Sultan, Inge MN Wortel, Jérémy Postat, Maryl Harris, Mauricio Merino, Judith N Mandl, Johannes Textor

## Abstract

Traditional cell tracking methods focus on fully automated tracking. The quality of the generated tracks, however, is heavily influenced by factors such as cell speed and density. In complex scenarios where track correctness is essential, an efficient and reliable tool for manual track correction remains a key component of the cell tracking process. Most tracking tools that allow for such manual correction do so only as a post-processing step, using human feedback to correct one track error at a time – without improving the quality of other tracks in the dataset. This highlights the need for a more efficient track correction method that iteratively integrates human feedback into the automated tracking process.

In this paper we propose a human-in-the-loop approach to cell tracking and investigate the extent to which a tracking algorithm can learn from human input *during* the tracking process. We combine a graph representation of cell detections, Dijkstra’s shortest path algorithm, and a neural network that predicts cell displacement to generate initial track suggestions. We introduce a tool that enables human feedback on individual tracks suggested by the algorithm; this feedback is then propagated to improve subsequent track generation. Compared to the traditional approach of fully automated tracking with correction as a post-processing step, our human-in-the-loop tracking framework reduces the number of corrections required. Finally, we show how this approach can be integrated with existing state-of-the-art tracking algorithms by using pre-generated tracks as a prior, improving the quality of tracks suggested to the human annotator.

**Author summary:** We introduce a novel human-in-the-loop method to improve efficiency in cell tracking, crucial for immune cell motility analysis. Automated tracking methods often struggle in complex environments, while manual corrections are more accurate but very time-consuming. Our method integrates iterative human feedback directly into the tracking process using graph-based cell detection, Dijkstra’s shortest path algorithm, and neural network predictions for cell displacement. Using this method as a backbone, we created an interactive tool that efficiently incorporates user corrections of individual tracks to progressively improve the overall track quality. Testing on T cells and neutrophils in microfluidic channels demonstrates significant reductions in required corrections compared to traditional methods, offering a scalable and efficient solution suitable for high-throughput biological research.

## Introduction

Cell migration is a crucial element of various physiological processes, such as tissue regeneration, immune response to pathogens, and the development of cancer [1]. Modern imaging techniques generate large amounts of microscopy data capturing cell movement in complex biological scenarios. This has naturally created a demand for high-throughput methods for extracting cell trajectories, also known as *cell tracking*. Currently, cell tracking algorithms can be grouped into two major categories. In the *tracking-by-detection* approach, one first segments (or directly finds the positions of) the cells, and then links the detections to tracks in a second step [2–5]. In the *end-to-end machine learning* approach, both object detection and tracking are done in a single step [6], often by using video foundation models [7]. This paper focuses on the tracking-by-detection approach, which is based on explicit assumptions about possible cell movements – as such assumptions can fundamentally influence the conclusions of downstream analyses, making these transparent can be required for valid analysis in many domains [8].

Tracking-by-detection consists of two steps. First, cells are segmented in each frame, leading to a set of *cell detections* (here defined as the coordinates of the centroid of the nucleus). Second, these detections are linked across time to form trajectories, or *cell tracks*, using one of several approaches: semi-automated, automated or manual. Manual tracking is time-consuming but more accurate, while automated approaches are scalable but more error-prone – especially in environments with a higher density of cells. For applications where having error-free tracks is essential – such as tracing cells over longer time periods to determine their long-term persistence – or the quality of automatically generated tracks is too poor, users either revert to laborious manual tracking or perform extensive manual correction of automatically generated tracks [9–11].

In some tracking tools, such as TrackMate [12, 13], manual correction is available as a post-processing step. A major limitation of this approach is that user feedback is only used to correct individual tracks and does not improve the quality of any other available tracks. Semi-automated cell tracking tools, such as ilastik [14] and ELEPHANT [15], require the user to provide some feedback, based on which a model is trained to generate tracks. While these approaches allow for iterative improvement, they are not explicitly designed to utilize the user’s feedback as efficiently as possible – for instance, an entire set of cell tracks is generated at once, which can make it time-consuming for users to find and repair potential errors.

In this paper, we set out to develop a human-in-the-loop (HITL) tracking approach that aims to minimize the amount of user feedback. In machine learning, “HITL” is an umbrella term that describes interactions between a user and a machine learning algorithm. We focused on one particular type: interactive learning [16], where the user provides iterative feedback and drives the system towards intended behavior [17]. Specifically, we propose an iterative HITL approach to cell tracking where we generate one track at a time, gather human input to correct it, and use this feedback to improve track generation later in the process, saving the user time and effort. Using simulations to estimate the amount of human work required for track correction, we show that our HITL method requires less human correction compared to baseline automated tracking methods combined with post-processing. We show that the number of required correction steps can be further reduced by improving the quality of the proposed tracks, either by incorporating the cell’s predicted displacement into the linking algorithm or by using the tracking output from baseline tracking algorithms as input. Finally, we present a manually curated dataset with 7629 tracks from 56 videos of T cells and neutrophils moving in channels of 2, 5, 6, 15 and 30 µm or a more complex 4-way split channel setup.

Overall, we find that our HITL approach, combined with cell-displacement prediction, as well as a prior of TrackPy tracks, reduces the number of track corrections required, compared to the baseline of manual correction of TrackMate tracks, for both 1D and 2D data.

## Materials and methods

### Data

The data that was used in this paper was generated for a study of T cell migration in microfabricated channels; we refer the reader to that study for details on the cells and imaging setup [9]. The data consisted of videos of neutrophils and T cells, moving in microfabricated channels of various widths and geometries (Supplementary Table S1).

The videos used were either in .czi or .tif format, which we read into Python using the czifile [18] and tifffile [19] libraries. We then converted each video into a series of 8-bit RGB frames using scikit-image [20], csbdeep [21], and numpy [22]. Each raw fluorescence channel was normalized independently by percentile contrast stretching, using csbdeep’s normalize function to clip intensities below the 3rd and above the 99.8th percentile before rescaling to the 0–255 range. We mapped the nuclear (DNA) channel to the red channel and the actin (LifeAct) channel to the green channel of the resulting RGB image, leaving the blue channel empty.

The majority of the obtained videos had dimensions 1376x1104x3, with pixel size 0.454×0.454 µm, or 2752×2208×3, with pixel size 0.227×0.227 µm. For convenience when training and testing our neural network, we reduced the size of the larger images to 1376×1104×3 to ensure all images had the same size and resolution.

### Cell position extraction using StarDist

The first step of tracking by detection involves finding the position of each cell in each frame, here defined as the center of the cell nucleus extracted from the (red) nucleus channel from the preprocessed RGB images (see above).

In order to extract the x- and y-coordinates of the nuclei we initially used a pretrained version of StarDist (`2D_versatile_fluo`; [23, 24]). For videos with lower density of cells, StarDist performed well in segmenting and outputting the cell nuclei centers. However, for videos with a higher density of cells or contaminated with cell debris, it output extra positions that did not correspond to actual cell nuclei. In addition, StarDist struggled with the accurate segmentation of neutrophil nuclei, often outputting multiple positions for a single nucleus. The reason for this is that the nuclei of neutrophils are segmented and have multiple lobes.

We developed a simple tool for adding missing detections and removing spurious detections from the StarDist output. The resulting set of corrected cell positions was then used for generating tracks and later for training our own neural network for cell segmentation and displacement prediction (see below).

### Generating cell tracks using existing methods

As a baseline for detection-based tracking, we used our manually curated detection sets (described above) and compared our tracking approach to two popular existing tools: the Track-Mate [12] plugin for Fiji [13] and the TrackPy [25] library in Python. Keeping the detection set the same allowed us to focus our comparison on the tracking algorithms themselves.

In TrackMate, we experimented with two of the available tracking methods: the Linear Assignment Problem (LAP) (**TrackMate-LAP**) and the Advanced Kalman Tracker (AKT) (**TrackMate-AKT**). For the latter, we empirically set both the initial search radius (used to link spots when a new track is started) and the search radius (used around the Kalman-predicted position for existing tracks) to 30px, and we set the maximum gap to 0 as a fair comparison to our algorithm, which does not allow gaps in tracks. To ensure reproducibility, we did not run these algorithms from the Fiji user interface but called TrackMate from Python using the TrackMate CSV Importer and the imagej and scyjava Python libraries. This allowed us to generate tracks for all videos for each tracking approach at once, instead of manually selecting the parameters for each video and each approach.

In TrackPy, we used either the basic cell tracking approach of TrackPy, the *link df* function (**TrackPy-Basic**) with a search range of 30px and a memory of 0, or the *NearestVelocityPredict* function (**TrackPy-NVP**), which attempts to improve the tracking by estimating and utilizing each tracked cell’s velocity (similarly to our own displacement prediction explained below). For TrackPy-NVP, we set the initial guess positions and velocities to None and used a span of 3.

For both the TrackMate and TrackPy approaches, we initially generated tracks only for individual channels of videos with narrow channels (2, 5, 6 µm). Later we tested our tracking approach on tracks from wider channels of 15 and 30 µm.

### Generating a detection graph

The approach to tracking that we chose is similar to that proposed by Magnusson et al. [26] and uses a graph representation of cell detections across frames, with nodes representing cell detections and edges representing possible connections between cell detections in consecutive frames. These edges are assigned a different “cost” based on some criterion deciding how likely these detections are to belong to the same cell (e.g. based on Euclidean distance or a more complex criterion, described further below). This detection graph is then used by the algorithm to generate tracks by finding low-cost paths through the graph.

To construct this detection graph, we distinguish five types of nodes, as defined in Table 1.

**Table 1:** Node types in the detection graph.

| Node type | Description |
| --- | --- |
| D | corresponds to a cell detection |
| Y | dummy node, used when a track does not start from frame 1 |
| X | dummy node, used when a track ends before the last frame |
| S | source node, used for finding shortest path |
| T | target node, used for finding shortest path |

Consider a video with *F* frames, indexed *t* = 1, …, *F*, where frame *t* contains *N*_*t*_ detections. The detection graph *G* = (*V, E*) has the node set

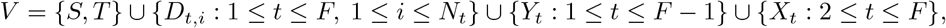

and the edge set *E* consists of the following edges with weights *w*:

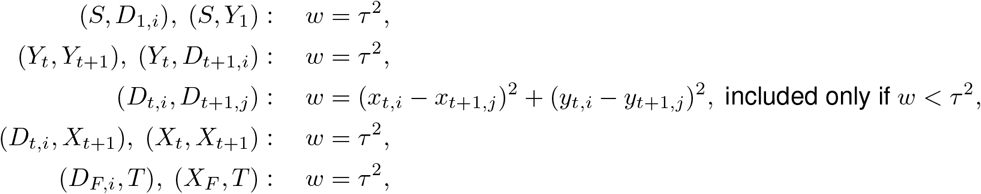

where (*x*_*t,i*_, *y*_*t,i*_) are the coordinates of *D*_*t,i*_, and *τ* is a predefined threshold representing the maximum plausible Euclidean distance that a cell can move between two time points.

The weights for the dummy SD-, SY-, YY-, YD-, DX-, XX-, XT-, and DT-edges were assigned a value of *τ* ^2^, the maximum squared distance allowed between two cells in two consecutive frames, which at the same time is also the value used to define which DD-edges to include. We also experimented with using different, higher weights for the SY-, YY- and DX-edges that influence where tracks begin and end, but this did not seem to have much of an effect (Supplementary Figure S1) so we decided to keep these weights all equal to *τ* ^2^ for simplicity.

### Cell tracking algorithm

Once the detection graph is constructed and edges have been assigned a cost, we used Dijkstra’s [27] shortest path algorithm to generate cell tracks one at a time. Intuitively, Dijkstra’s algorithm is a reasonable choice for our problem because for a detection *D*_*m,n*_ in frame *m*, we would assume that the detection in frame *m* + 1 with the smallest cost from *D*_*m,n*_ would be the realization to the same cell. However, this assumption may not hold if cells move fast enough to displace by much more than their own size per frame.

Importantly, after a track is generated by finding the shortest path between the source node S and the target node T, the D-nodes of this path can be removed from the detection graph as they no longer need to be considered for future tracks – gradually simplifying the problem as more tracks are accepted. The process was completed when no D-nodes were left in the graph.

While for some cell videos this algorithm produced mostly correct tracks, for others it performed worse, generating tracks with many errors.

These varying results suggested that we needed an improved cost function which would take into account the displacement of a cell between two frames.

### Using candidate tracks as a soft prior

In some experiments, we already have a set of candidate tracks available from an existing tracking algorithm (e.g., TrackPy). To bias our algorithm towards generating tracks from this input set whilst still allowing it to overrule the input if necessary, we lower the cost of edges corresponding to the input tracks. In this paper, we implemented this “soft prior” by dividing each edge weight corresponding to an input track by *τ* ^2^, giving such edges a cost *≤* 1.

### Artificial neural network for improving the cost function

#### Improved cost function

Instead of using the squared Euclidean distance as a cost function, we trained a neural network to predict the displacement on the x- and y-axis of each cell between two consecutive frames, with the new cost between two detections *D*_*t,i*_ and *D*_*t*+1,*j*_ becoming

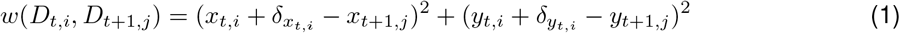

where 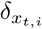 and 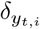 represent the predicted x- and y-displacements of *D*_*t,i*_ between time points *t* and *t* + 1. Instead of favoring detections *D*_*t*+1,*j*_ with minimum distance to *D*_*t,i*_, this favors detections with minimum distance to the *predicted* position 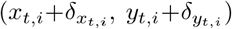. We established empirically that the threshold for the Euclidean distance of 30px was still a good choice.

### Choice of neural network architecture

For the task of improving our cost function by predicting cell displacements, we used a U-Net. U-Net is a type of neural network which is widely used for cell segmentation [28]. It consists of a contracting path to capture features and context of an image and an expanding path for localization and reconstruction. In our case, our main goal was to predict the displacement of cells in the x- and y-direction in the next frame, but we also used the network for cell segmentation. We decided to also train the network to segment cells because this is extra information which could potentially also improve the displacement predictions. In addition, given our encoding of the displacement as color intensity, a non-displacing cell is encoded in the same way as no cell at all, so without the additional segmentation channel we would be unable to detect non-moving cells.

### Training input and output

RGB images extracted from cell videos were split into non-overlapping square patches of size 128×128×3. This was necessary as the neural network’s performance was poorer when using the entire images as an input. The input to the neural network consists of two stacked RGB images, corresponding to the same area of two consecutive video frames, with size 128×128×6. We wanted our neural network to generate as output all elements that we required for cell tracking – cell positions and predicted displacement of each cell in x- and y-direction in the following frame. For each image corresponding to a video frame, we generated 3 images that were then used in the training output. We generated a grayscale image with a black background and white disks with radius r=10 pixels, centered on the nucleus centers extracted using the manually cleaned detection set (see above). We also used the tracks we had already generated using our tracking tool to extract the x- and y-displacement of each cell, using the difference between the next frame’s and the current frame’s x- and y-coordinates.

To encode these ground-truth displacements from a pair of frames, for each cell we generated two separate images, one encoding the x-displacement and the other the y-displacement. We used two channels per dimension: one to encode positive displacements, and the other to encode negative displacements. The final training target used was the signed difference between both channels. This setup allowed us to easily visually investigate the training data and network predictions. For each cell, we filled a disk with a 10-pixel radius centered at the cell’s first position with the displacement encoded as described. Displacement predictions were then made by averaging the network-generated image over the 10-pixel-radius disk centered at the cell’s detected position.

The neural network training output thus consisted of three channels: one corresponding to the cell positions, one corresponding to the x-displacement, formed by taking the difference between the positive- and negative-displacement channels, and one for the y-displacement, formed in the same way. In the same way as the input, these were split into patches, making the size of the output 128×128×3.

### Neural network implementation and training

For training the neural network we used PyTorch version 2.1.0.dev20230525. The final network used in this paper was trained on 66,905 128×128×6 input tiles extracted from consecutive frame pairs of the 17 training videos (Supplementary Table S1). The network was trained to minimize the mean squared error between the predicted and target output channels (segmentation mask and x-/y-displacement maps). We used the Adam optimizer [29] with a learning rate of 0.0001. We used L1 regularization with weight decay *λ*_1_=0.1 and L2 regularization with *λ*_2_=0.0001. In addition, we applied data augmentation to the input. The data augmentation of each image consisted of applying a horizontal flip, vertical flip, adding Gaussian noise with mean 0 and standard deviation 0.05, and adjusting the brightness with a brightness factor drawn from a uniform distribution on [1, 3]. Each of these augmentations was applied with a probability of 0.5 during training.

Training was run for 159 epochs, which took about 22 hours on a MacBook Pro (2020), equipped with M1 chip, 8-core CPU and 8-core GPU. To leverage GPU capabilities, the PyTorch Metal back- end was used. We used scikit-learn [30], scikit-image [20], and numpy [22] for some important pre- and post-processing of the input and output.

The neural network used to generate displacement predictions for this manuscript was trained using an initial version of the tracks where some, but not all, were double-checked and manually corrected for any residual errors. The reason for this was that the final, fully curated dataset was obtained using the tool with the displacement prediction network. While this may slightly affect the performance of the displacement network itself, we do not expect it to have a major effect on the performance of the full HITL pipeline.

### User interface for enabling human input for our online algorithm

While we use Dijkstra’s shortest path algorithm for suggesting cell tracks, the user interface we developed [31] enables human input, effectively turning our tracking approach into an online algorithm.

#### Backend

On the backend our user interface generates a detection graph *G* based on a provided CSV file with detections. It then uses an implementation of Dijkstra’s shortest path algorithm to generate a single track proposal. This proposal is then displayed, using the frontend, where the user can inspect the track proposal. The actions that the user can take can be seen in Table 2. Each action changes the detection graph *G* by adding or removing nodes or edges. Once the changes are made, a new track proposal is generated using Dijkstra’s shortest path algorithm. This track proposal can be of a different track, as the actions taken by the user can affect the detection graph in such a way that the previously proposed track no longer corresponds to the shortest path. The backend of our user interface was developed entirely in Python.

**Table 2:** Comparison of actions for track correction in our tool (HITL) and TrackMate (TM)

| Action | Key combinations |  | Number of clicks |  | Simulation actions |
| --- | --- | --- | --- | --- | --- |
|  | HITL | TM | HITL | TM |  |
| Remove detection | **D** (for currently selected detection) | **Left-click mouse + D** | 1 | 2 | 1 |
|  | **Right-click mouse** (on any detection) |  | 1 |  |  |
| Remove edge | **Left-click mouse** on correct point + **W** | **Left-click mouse** on edge, then **right-click** → *Remove links only* | 2 | 3 | 1 |
|  | **X** to delete all outgoing edges from current detection. |  | 1 |  |  |
| Remove track (segment) | **Right arrow** $n - 1$ times to inspect track of length $n$, then press **Shift + Z** | **Down arrow** $n - 1$ times, then **right-click** → *Select track upwards*, then **right-click** → *Remove spots and links* | $n + 1$ | $n + 3$ | 1 |
| Add detection | **Shift + left-click mouse** | Hover mouse to correct position + **A** | 2 | 1 | 1 |
| Add edge | **Left-click mouse** on first point, then second point (next frame), then press **W** | **Left-click** on first point, drag edge to second point, release | 3 | 1 | 1 |
| Add consecutive edges | **Left-click mouse** on each of the $n$ detections, then press **W** | **Shift+Left-click** each of $n$ detections, then **right-click mouse** → *Link $N$ Spots*<br>Drag to select all spots, then **right-click** → *Link $N$ Spots* | $n + 1$ | $n + 2$ | $n - 1$ |

#### Frontend

The frontend of our interface was developed using napari [32]. The interface displays one frame at a time, controlled by the user through the left and right arrow keys. It also shows the track generated on the backend, with the links between nodes displayed as white lines, as well as the current position of the track in the frame at hand.

The user can then make corrections or accept the track, using the operations in Table 2. When the graph is updated, accounting for the corrections, a new track is generated, again using the shortest path algorithm on the reduced graph.

### Metrics for comparison of tracking methods

In order to compare the tracking methods, we used the Track Fraction (TF) and Complete Tracks (CT) metrics as defined in [33]. In this context, a track fraction is the longest continuous correctly reconstructed fraction of a track. The TF metric is then calculated by averaging all track fractions in a video. The CT metric is calculated as

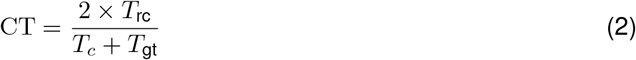

where *T*_rc_ is the number of ground-truth tracks that were completely reconstructed by the algorithm, *T*_gt_ is the number of all ground-truth tracks, and *T*_*c*_ is the number of tracks computed by the algorithm.

## Results

### A manually curated *in vitro* dataset for evaluation of cell tracking pipelines

As a basis for the work presented in this paper, we made use of a large dataset that we recently generated [9], in which we imaged T cells and neutrophils *in vitro* in microfluidic devices. Specifically, we acquired 56 videos in which the cells were confined to channels of varying widths (ranging from 2–30 µm; Supplementary Table S1).

This microchannel dataset had several important characteristics that were relevant for our work. First, while we generated initial candidate tracks using a combination of open-source tools (similar to the ZeroCostDL4mic approach [34]; Methods), tracking in this scenario was sufficiently challenging that we had to manually correct each track in a laborious but necessary process, which motivated this research. Second, the channels varied in the number of cells they contained, meaning that the data represented varying degrees of difficulty. Third, in a large fraction of the videos (T cells in at most 6 µm wide channels), cells were unable to pass each other, effectively constraining the motion to one dimension. This made it possible to manually identify a high-confidence “ground truth” in many of the videos, even when these were not necessarily easy to track by existing methods. Finally, the dataset included several videos of migrating neutrophils. Neutrophils are especially challenging to track due to their ability to hypersegment their nuclei (“pearl-on-a-string configuration”), which helps them to move in confined environments [35]. This unusual nucleus configuration can throw off pre-trained cell detection networks such as StarDist, which makes these cells especially challenging to track and therefore an interesting candidate for HITL tracking. On the other hand, their ability to pass each other even in narrow microchannels means that the ground truth for these cells is also somewhat less certain in some cases when many cells cluster together.

Overall, we collected a dataset of 7629 tracks representing two different types of cells moving in various *in vitro* environments at varying speeds and densities. We used *∼* 70% of these data to test our approach in the simulations we developed, where test data was chosen to represent various scenarios of cell movement – from low-density videos with many individual cells to high-density videos containing many cells moving collectively (Supplementary Table S1).

### A human-in-the-loop tracking algorithm that incorporates optional user feed-back in the tracking process

Having witnessed the amount of work required to construct large manually curated cell tracking datasets as a foundation for scientific analysis, we set out to develop a more efficient approach where the inevitable corrections made by the human operator are leveraged to improve their experience going forward. We therefore developed a HITL cell tracking approach which includes user feedback as part of the tracking process, instead of as a post-processing step.

This method has three steps: cell segmentation, detection graph construction, and HITL tracking. The segmentation step can be performed either by a standard cell segmentation model, such as StarDist, or using the neural network that we proposed for cell segmentation and displacement prediction for our specific dataset (see Methods).

The extracted cell detections (Figure 1A) are then used to generate a detection graph with nodes corresponding to the detections and edges between cells in consecutive time points. These edges are assigned a weight that reflects how likely these detections are to belong to the same cell according to the chosen cost function (either the squared Euclidean distance or based on predicted displacements, see Methods for details). Finally, tracks are generated using Dijkstra’s shortest path algorithm to find tracks with a low cost (Figure 1B). In our particular implementation, the graph algorithm runs as a backend and generates one track at a time, which is then presented to the user for correction through a frontend. The user can then either correct a track error or accept a completely correct track. The feedback at every iteration is used to update the graph and simplify it, which automatically improves the quality of tracks suggested later in the process.

**Figure 1.**
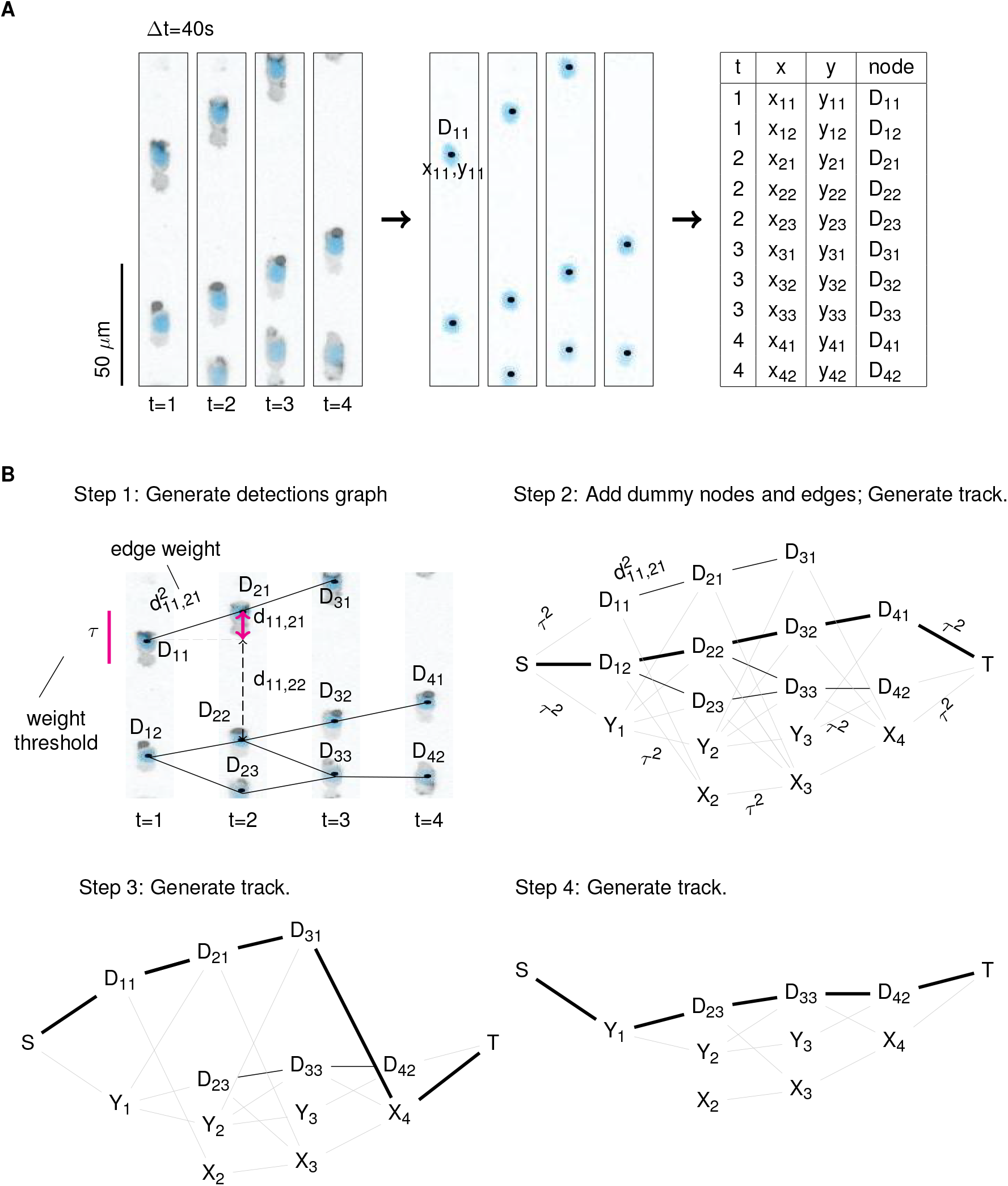
Cell segmentation, detection graph generation and tracking. **(A)** The coordinates of nuclei centers are extracted, using StarDist or another detection network. **(B)** The coordinates are used to generate a detection graph. Cell tracks are generated by iteratively finding the shortest path between the source node S and the target node T, using Dijkstra’s shortest path algorithm, removing the D-nodes included in a shortest path. Cell tracking is complete when no detection nodes are left in the graph.

In summary, our HITL tracking approach – implemented as a napari plugin [32] – iteratively generates one track at a time and allows human feedback, which is then used to indirectly improve the quality of any further generated tracks.

### A simulation setup to estimate the amount of work needed for cell tracking

We developed a simulation to investigate the effect of individual pieces of feedback on the tracking process, evaluating the change in the average track fraction, proportion of complete tracks, and the number of accepted tracks on the complete set of tracks generated after each step.

At each iteration a single track is generated and matched to the ground-truth track of its very first detection node. The two are then compared and a single user action is simulated: for example, accepting the whole track if it is fully correct, or removing or adding an edge if it is not. The user feedback is then used to modify the detection graph. Then, the effect of that change is evaluated by automatically generating all remaining tracks from the updated graph (together with any that were already accepted), and compared to the ground truth in terms of TF, CT, and the number of accepted tracks. The process is then repeated with a new suggested track until all suggestions have been presented to the user for feedback and the detection graph is empty.

In order to evaluate the work required to obtain a completely correct set of tracks, we evaluated the proportion of correction iterations *p*_corr_, i.e., the ratio (# correction operations)/(# correction operations + # track-accepted operations).

As can be seen in Figure 2, the values of TF (Figure 2A) and CT (Figure 2B) increase over the course of the tracking process, but can temporarily decrease after accepting a new track (Figure 2C). This is due to the fact that each detection usually has several outgoing edges. Thus, a single correction introduced to the detection graph could then force the algorithm to choose different paths when re-generating a complete set of tracks on the same detections, resulting in some new track errors. These are however temporary, with the HITL tracking process resulting in a clear increase of track quality over the iterations. There is also a clear inverse correlation between the amount of work required to correct tracks and the initial quality (Figure 2D: TF; Figure 2E: CT).

**Figure 2.**
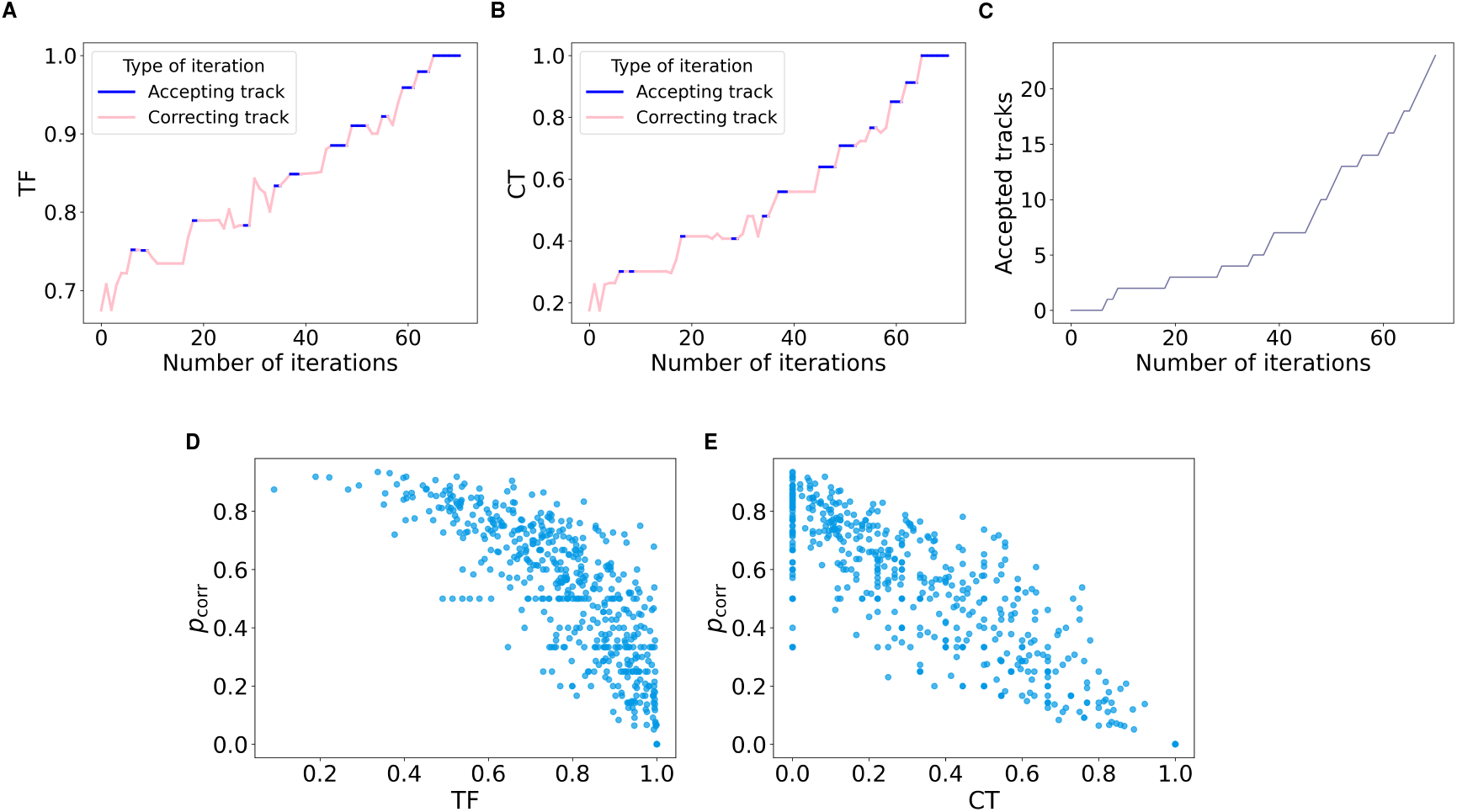
A simulation of our human-in-the-loop approach helps evaluate the amount of work required for correcting a set of cell tracks. **(A-C)** Measuring the quality of tracks after each iteration (A: TF; B: CT; C: number of accepted tracks over time for an example dataset) helps evaluate the amount of work needed for track correction. **(D-E)** A higher quality of the initial set of tracks (D: TF; E: CT) is associated with fewer correction iterations (Y axes). Each point stands for one channel.

Based on these results, we decided to test two further approaches to reducing the amount of required correction work: incorporating additional information into our own tracking algorithm, and using alternative, state-of-the-art tracking systems as prior input.

### Displacement prediction further reduces the amount of required feedback

When testing our HITL tracking tool, we noticed that many errors were due to cells moving about their own size in a single frame and occupying the same position where an earlier cell had been in an earlier frame. This meant that the common approach of connecting the detections that are closest to each other was not performing well. To improve the quality of the initial tracks suggested by our algorithm, we therefore investigated the effect of adding displacement prediction to the cost function.

Specifically, we trained a neural network to predict the x- and y-displacement of cells in the following frame (Figure 3). The network is trained on pairs of frames, and then predicts a displacement vector for each identified cell in a single frame at inference time. The paired frames allow the network to take e.g. cell morphology and actin polymerization into account when it predicts the most likely next position. We then modified our cost function (Equation 1) as follows: when two frames at times *t* and *t* + 1 are compared, we substitute the positions of the cells in frame *t* by their predicted positions at time *t* + 1.

**Figure 3.**
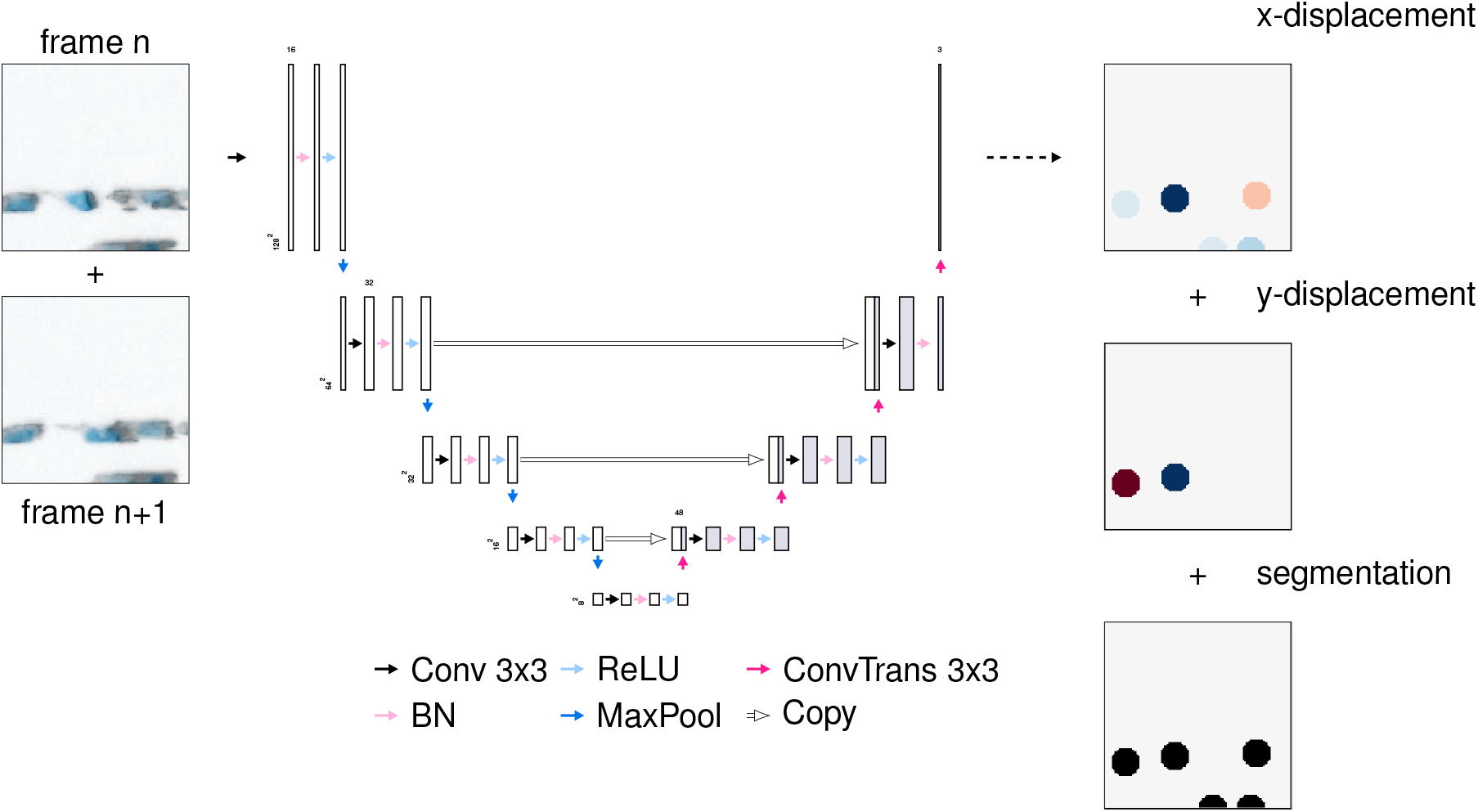
U-Net architecture, input and output. The neural network takes as an input two stacked consecutive image tiles (128×128×3) and outputs a segmentation mask, as well as displacement masks for the x-axis and y-axis displacement.

We then evaluated the benefit of the displacement predictions by iteratively running Dijkstra’s algorithm to generate shortest paths, with no human feedback. We found that adding the displacement prediction overall increased the quality of the initial tracks used for human input, and that it also made the algorithm less sensitive to the choice of the maximum distance threshold (Table 3).

**Table 3:** Adding displacement prediction to the cost function improves baseline track quality. Based on these results, we set the cutoff threshold for the displacement-modified Euclidean distance to 30px.

|  | T cells |  | Neutrophils |  |
| --- | --- | --- | --- | --- |
| without displacement | TF | CT | TF | CT |
| $\tau = 15$ | .723 (.775, .547-.963) | .275 (.154, .007-.41) | .505 (.474, .276-.717) | .096 (.015, 0-.07) |
| $\tau = 20$ | .921 (.987, .904-1) | .552 (.567, .2-1) | .645 (.66, .439-.864) | .198 (.062, 0-.221) |
| $\tau = 25$ | **.941** (.993, .912-1) | .63 (.667, .316-1) | .8 (.853, .688-.978) | .32 (.193, .056-.493) |
| $\tau = 30$ | .936 (.987, .901-1) | **.642** (.696, .308-1) | .888 (.924, .811-1) | .463 (.369, .135-.95) |
| $\tau = 35$ | .929 (.983, .893-1) | .637 (.667, .296-1) | **.896** (.931, .817-1) | **.539** (.5, .224-1) |
| with displacement |  |  |  |  |
| $\tau = 15$ | .964 (1, .975-1) | .762 (.96, .581-1) | .907 (.948, .833-1) | .524 (.471, .202-1) |
| $\tau = 20$ | **.968** (1, .974-1) | .805 (1, .667-1) | **.915** (.958, .844-1) | .588 (.571, .298-1) |
| $\tau = 25$ | .965 (1, .97-1) | .81 (1, .667-1) | **.915** (.97, .845-1) | .626 (.622, .333-1) |
| $\tau = 30$ | .959 (1, .961-1) | **.812** (1, .667-1) | .904 (.965, .829-1) | .644 (.633, .351-1) |
| $\tau = 35$ | .955 (1, .952-1) | .801 (1, .636-1) | .897 (.957, .816-1) | **.656** (.667, .383-1) |

In summary, we found that a simple displacement prediction generated by a neural network improved the initial quality of the tracks. This illustrated the flexibility of our HITL approach to leverage machine learning analysis of the underlying videos.

### Incorporating a state-of-the-art tracking algorithm as a prior improves human-in-the-loop tracking

While repeated execution of Dijkstra’s algorithm is an approach that lends itself particularly well to a HITL setup as it generates tracks one-by-one, this kind of greedy approach may not be the best overall. We therefore also investigated whether initializing the approach with tracks generated using a different, independent algorithm would reduce the number of required corrections.

To incorporate this kind of “prior” input from an independent algorithm *A* into our approach, we modified the initial graph construction. In a “soft prior” approach (Figure 4A), we lower the cost of edges corresponding to tracks generated by algorithm *A*. This means that Dijkstra’s algorithm will generally prefer to generate such tracks, but may choose to diverge from the input if it finds a much better option. The “soft prior” approach will be used to determine if using a better tracking algorithm as input reduces the required correction work.

**Figure 4.**
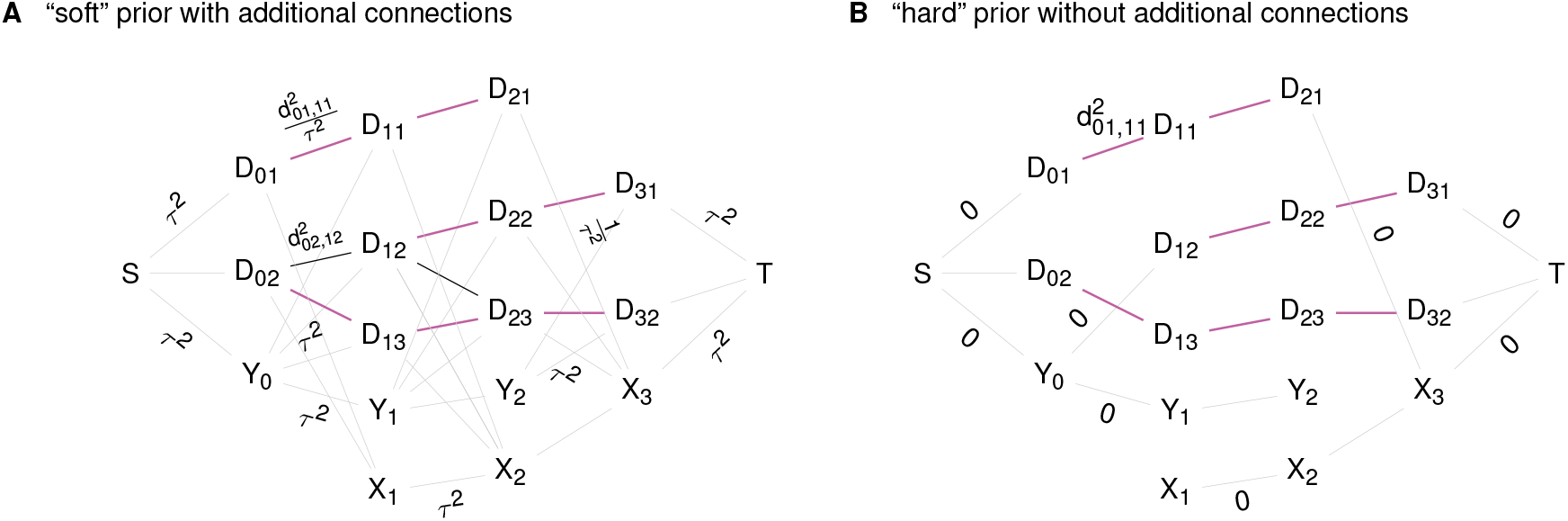
Initialization of the tracking graph with tracks generated by another algorithm. **(A)** In one approach, we initialize the detection graph using all tracks found by the algorithm (purple edges) with suitably reduced cost (e.g., by dividing the cost by *τ* ^2^) and then add potential connections (of cost *≤ τ* ^2^) between detections in adjacent frames (gray edges). **(B)** Alternatively, we add only the tracks found by the algorithm without additional edges. These would then have to be added back by the human operator.

Alternatively, in a “hard prior” approach (Figure 4B), we only have edges corresponding to the initially generated tracks, which means there is no way for Dijkstra’s algorithm to diverge from this input except if the user makes corrections. We will use this “hard prior” approach to quantify how much work it takes to correct the tracks produced by one of the baseline algorithms.

We chose two baselines for our experimentation: the software “TrackMate” [12], which is very popular in the cell migration field, and the Python package “TrackPy” [25], which was easy to integrate into our tooling. Specifically for TrackMate, we also developed a simulation of the manual correction tool offered by the software itself, which uses different operations than our own (Table 2, Methods). When used on our data, all algorithms showed a decent to good baseline performance (Table 4).

**Table 4:**
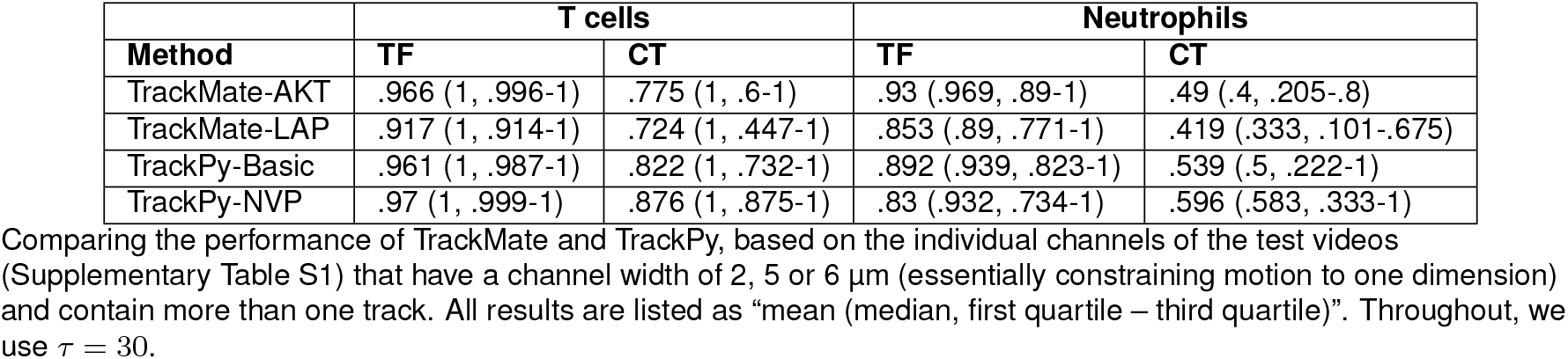
Comparing the performance of TrackMate and TrackPy on 1D tracks.

Encouraged by these results, we next performed track correction simulations starting from these different baselines. Since the cell densities vary across channels and videos, we performed this evaluation separately on *simple* and *complex* channels. Specifically, we defined a simple channel as a channel where fewer than 25% of the cells from all frames are closer than 30px to at least one cell in the same frame; other channels were considered to be “complex”. In both simple (Figure 5A) and complex channels (Figure 5B), combining TrackPy tracks with our method and displacement prediction reduced the number of corrections required, compared to the baseline of TrackMate tracks corrected with TrackMate’s manual tool. Overall, the best performance was achieved when using TrackPy as a soft prior and leveraging our displacement predictions.

**Figure 5.**
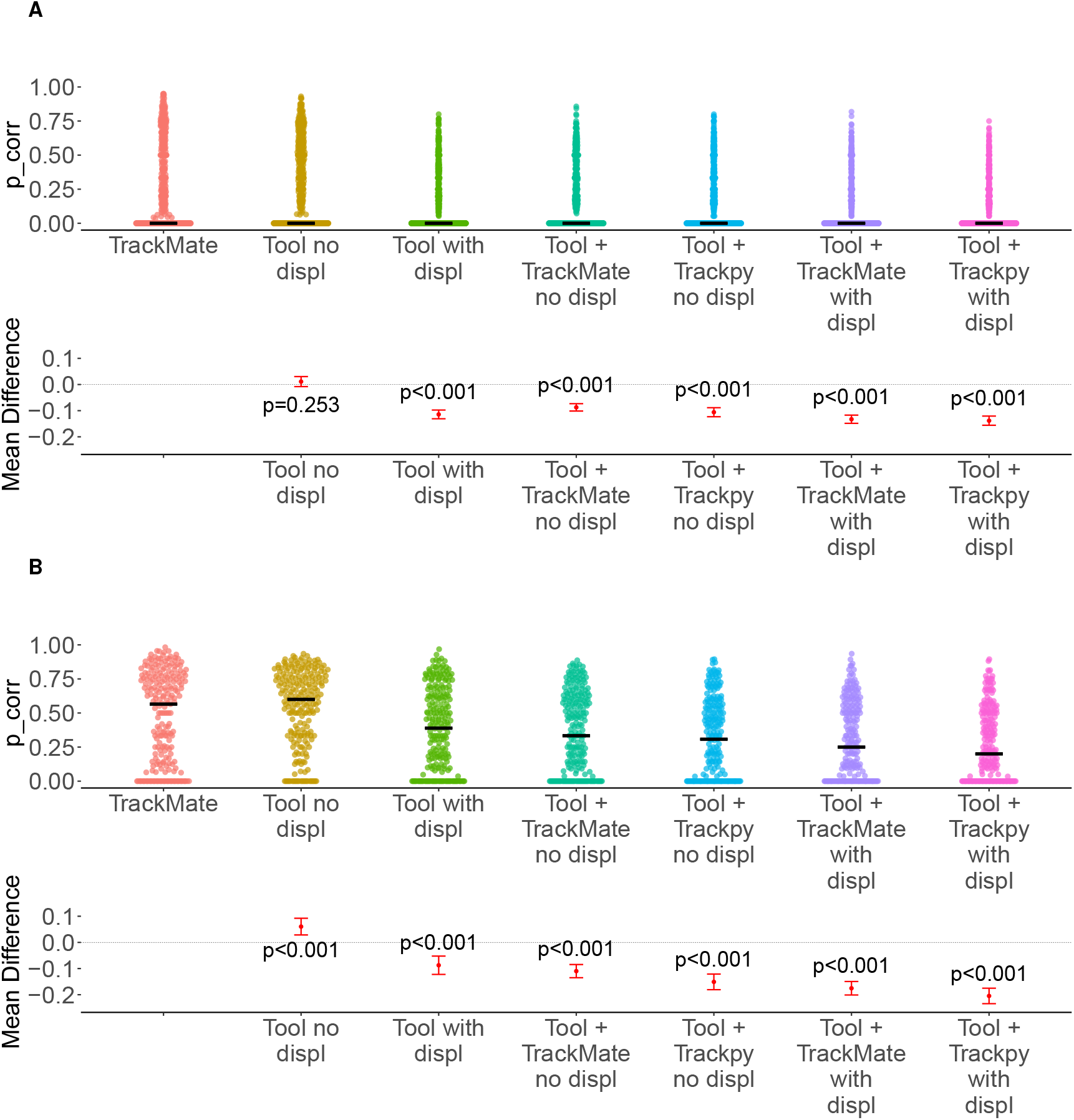
Starting with better initial tracks decreases the number of required corrections. Based on “simple” channels, where fewer than 25% of the cells from all frames are closer than 30px to at least one cell in the same frame. **(B)** Based on “complex” channels, where the percentage of “close” cells is more than or equal to 25%. Throughout, “TrackMate” uses the tracks from TrackMate as a hard prior; “Tool no displ” stands for the simple HITL algorithm without displacement prediction; “Tool with displ” is the same method with displacement prediction; and “Tool + TrackMate” and “Tool + TrackPy” stand for using the mentioned baseline algorithm as a soft prior. Top panels: proportion of correction iterations (*p*_corr_) per channel (points); black bars: medians. Bottom panels: mean paired difference in *p*_corr_ relative to “TrackMate” with 95% confidence intervals. The *p*-values are from two-sided paired *t*-tests without correction for multiple comparisons.

### Comparing tracking approaches for different target audiences on more challenging data

Having performed all experiments so far on the same large dataset, we asked to what extent the observed performance differences would translate into a reduced correction workload on a more challenging, real-world task. For these experiments we only used channels from two more challenging videos – one with 15 µm and one with 30 µm channels. Cells in these channels are harder to track since they can pass each other and generate complex “swirling” motion that is not observed in single-lane traffic.

We were interested in how big the differences between some of our setups would be in these more challenging data, and chose three setups that reflect three typical groups of users. First, we considered users who are most comfortable with using a completely off-the-shelf tool which is easy to install, such as TrackMate. These users would not be interested in installing something like our napari plugin for HITL cell tracking, and would use TrackMate itself to edit and fix any errors. Second, we considered users who would be comfortable using an off-the-shelf tool like our napari plugin, but may not have machine learning expertise and may not want or be able to train their own neural networks to generate displacement predictions. Third, highly technical users might simply use our provided Python library for the HITL tracking – perhaps writing their own frontend – and would be fine with training their own displacement prediction models. We considered such users to be most likely to use TrackPy rather than TrackMate. To ensure that the differences between the three approaches are due only to tracking, we used the ground-truth cell segmentations for these channels in all three simulations.

Our results show that in both simple (Figure 6A) and complex channels (Figure 6B) using TrackPy within our tool and displacement prediction (corresponding to a highly technical target audience) reduced the required number of corrections, compared to a baseline of TrackMate tracks corrected with its own manual correction tool. The combination of HITL track editing with a TrackMate prior performed quite similarly, but did not differ significantly from the TrackMate-only baseline.

**Figure 6.**
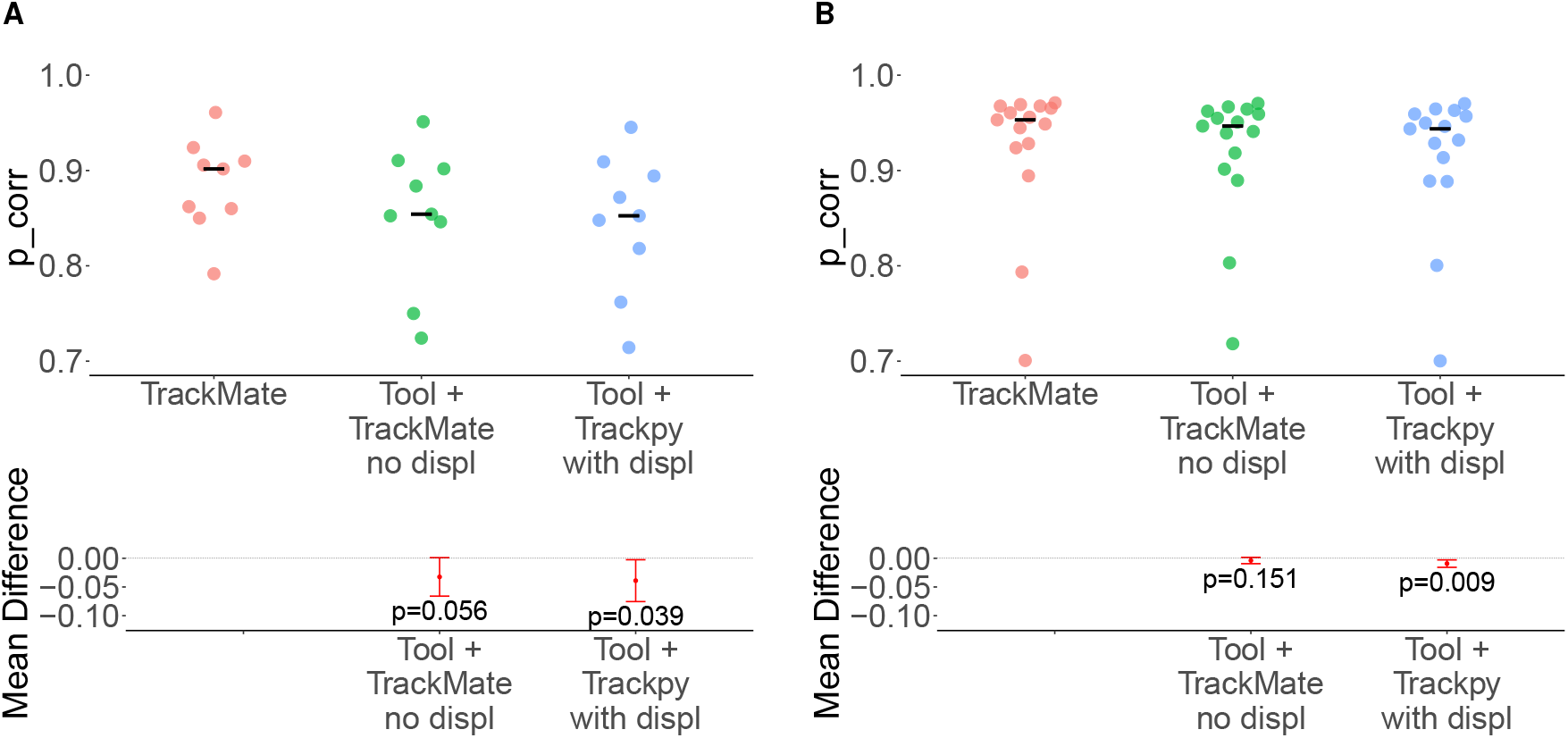
Comparing three typical approaches on wider channel data. **(A)** Simple channels; Complex channels. “TrackMate” corresponds to TrackMate tracks corrected with TrackMate’s own manual correction tool; “Tool + TrackMate no displ” uses TrackMate tracks as a soft prior in our HITL tool without displacement prediction; “Tool + TrackPy with displ” uses TrackPy tracks as a soft prior together with displacement prediction. Top panels: proportion of correction iterations (*p*_corr_) per channel (points); black bars indicate medians. Bottom panels: mean paired difference in *p*_corr_ relative to “TrackMate” with 95% confidence intervals; The *p*-values are from two-sided paired *t*-tests.

In summary, our experiments showed that a HITL approach produced noticeable reductions of work in a realistic setting, although we cannot conclusively state whether an improved cost function that incorporates cell displacement predictions is required for these improvements to materialize.

## Discussion

In this paper we proposed an iterative HITL method which incorporates track correction in the tracking process, instead of in post-processing. Traditional tracking tools, such as the popular Fiji plugin TrackMate, offer both automated and manual cell tracking. Manual tracking, while ensuring in theory error-free tracks, is very time-consuming and not scalable for high data throughput. Automated tracking produces high-quality tracks in some scenarios, but is still error-prone in others – for example, in our broad-channel datasets, very high numbers of corrections were still required. The notion that tracking is not a “solved problem” is also supported by the many novel tracking approaches that continue to be developed and published [4–6], in part motivated by a dissatisfaction with the performance of existing tools [10]. Some popular tracking tools, such as TrackPy, do not provide tools for manually correcting such errors at all; others, such as TrackMate, allow users to correct tracks as a post-processing step, where corrections are only applied to the track at hand and do not propagate to any other tracks. All in all, this illustrates the need for not just better automated tracking algorithms, but also better correction tools that combine the scalability of automated tracking with the precision of manual tracking.

We found that fewer corrections are required after incorporating TrackMate and TrackPy tracks into our method as a soft prior, as opposed to the baseline of manually correcting TrackMate tracks through post-processing only. This demonstrates that propagating feedback to any further generated tracks is more efficient than only applying corrections to the track at hand. Furthermore, this makes our tool compatible with any state-of-the-art tracking method which can generate a complete set of tracks. Predicting cell displacement between consecutive time points further improves the performance of our tool compared to the baseline, both with and without using a prior of TrackMate and TrackPy tracks. This shows that cell displacement is a predictable and useful element in cell tracking.

The idea of using some form of a HITL approach in cell tracking is not new. The two tools that are most similar to our approach are the tracking module of ilastik and the 3D-tracking tool ELEPHANT. Ilastik offers an iterative “tracking with learning” workflow. In this workflow, the user annotates a few short tracklets which are then used to generate weights for a probabilistic graphical model which is constructed for all detections in all time points simultaneously. After the model has been trained and run, the user can inspect the output tracks and, if needed, annotate some additional tracklets and retrain the model in order to improve the tracking output [14]. A similar approach has been used in ELEPHANT. The user links the detected cells in one frame to their corresponding detections in the next frame. Once again, sparse annotations are used. An optical flow model is trained, which predicts the displacement of the cells in the previous frame. Cell positions together with the predicted displacement are then used to generate cell tracks. The corrections of tracks are then used as labels to retrain the optical flow and improve displacement prediction [15]. In these tools a HITL approach has been used to generate model weights and improve the quality of the input of the tracking algorithm, respectively, which as a result improves the generated tracks. Both of these tools output a whole set of generated tracks, as opposed to our tool which presents the user with a single track at a time. Inspecting individual tracks makes it easier to identify errors and correct them. Furthermore, it allows the user to determine the amount of work they are willing to put into track correction. The user can correct tracks and also accept tracks that are completely correct. Thus, if they have a target number of completely correct tracks they want to generate, they can choose to stop as soon as this number is reached.

Despite incorporating human feedback during track corrections, the amount of work needed in our approach still depends on the performance of the tracking algorithm that is used to generate the initial and improved tracks. Tracking dense cell populations, for example, is a known problem for state-of-the-art algorithms: the proximity of cells makes the algorithm more error-prone, as there are more potential candidates to link a cell to [10]. Similarly, tracking cells in open 2D (or even 3D) environments is inherently harder, as there are more possibilities for the direction in which a cell would move in the next frame. This should also make training a robust model to predict cell displacements more challenging and data-intensive. It would be worth exploring how to combine cell movement simulations and generative machine learning models to create realistic videos of moving cells. These could then be used for training a neural network for displacement prediction in cases with scarce image data.

Another direction for future research would be to investigate whether there is a more effective way to use human feedback. Our current approach is based on using user feedback to simplify the detection graph. It would be interesting to combine this approach with a method similar to ilastik or ELEPHANT, where corrections are used to refine a machine learning model – in our case, this could mean that displacement predictions could be improved, leading to more useful edge weights in the graph and potential tracking improvements. Similarly, we have not explored in detail what the best way would be to use initial tracks as a “soft prior”.

In this work, we have used simulation to determine the amount of correction work, and measured the number of required operations. In reality, different operations will take different amounts of time, and users will need a while to find positions where corrections are needed in the first place. We believe that going systematically through the data track-by-track, focusing on high-fidelity ones first, may in fact reduce this search time. To properly evaluate the impact on user experience, usability studies with human users would be necessary.

In summary, our work introduces a HITL tracking framework which effectively incorporates user feedback into the tracking process. It offers an efficient alternative to manual track correction, with clear benefits for narrow-channel cell data. The possibility to integrate tracks generated by state-of-the-art tools such as TrackMate and TrackPy makes it a valuable addition to the computational biology toolkit.

## List of abbreviations

AKT: Advanced Kalman Tracker
CT: Complete Tracks
LAP: Linear Assignment Problem
TF: Track Fraction
HITL: human-in-the-loop

## Acknowledgments

We would like to acknowledge Lucas Wiebke, Daniel Parisi, and Shane Pullens for their help generating cell tracks, Daria Kondrateva for checking and correcting cell tracks, and Koert Schreurs for correcting cell tracks and improving the image processing pipeline. We also thank Lin Wouters, Kumaran Viswanathan, and David Cicchetti for their help improving the image processing pipeline. Finally, we thank Sara Ben-Hammou for her contribution to curating the final version of the dataset.

## Author contributions

- **Mihaela Mihaylova** — Conceptualization (equal), Methodology (equal), Software (lead), Validation (lead), Formal analysis (lead), Investigation (lead), Data curation (lead), Writing — original draft (lead), Writing — review & editing (lead), Visualization (lead).
- **Ankur Ankan** — Software (supporting), Writing — review & editing (supporting).
- **Shabaz Sultan** — Methodology (supporting), Software (supporting), Writing — review & editing (supporting).
- **Inge MN Wortel** — Conceptualization (supporting), Methodology (supporting), Writing — review & editing (supporting), Supervision (supporting).
- **Jérémy Postat** — Resources (equal), Data curation (equal), Writing — review & editing (supporting).
- **Maryl Harris** — Resources (equal), Data curation (equal), Writing — review & editing (supporting).
- **Mauricio Merino** — Resources (equal), Data curation (equal), Writing — review & editing (supporting).
- **Judith N Mandl** — Resources (equal), Writing — review & editing (supporting), Funding acquisition (equal).
- **Johannes Textor** — Conceptualization (equal), Methodology (equal), Data curation (supporting), Writing — original draft (supporting), Writing — review & editing (supporting), Visualization (supporting), Supervision (lead), Project administration (lead), Funding acquisition (equal).

## AI use disclosure

The authors used Claude Opus 4.8 and 5.5 (Anthropic), accessed through Claude Code, to proof-read manuscript versions for typos, grammatical errors, and logical gaps or inconsistencies. Each change suggested by Claude Opus was individually reviewed and modified as necessary. Claude Opus was not used to perform analyses, draw scientific conclusions, or draft any text beyond suggested modifications addressing specific flagged issues. In addition, ChatGPT-4-family models (OpenAI) and Claude Sonnet 5 and 5.5 (Anthropic) were used for coding assistance, to debug and improve code. All AI-assisted code was reviewed, tested, and validated by the authors. The authors take full responsibility for the content of this manuscript.

## Data and Code Availability

The raw image data and the manually curated tracks are available on Zenodo [36]. Our HITL tracking tool is available as a Python package called `trackin` on PyPI [37]. All code used to generate the figures in this paper will be shared publicly upon acceptance of this manuscript for publication, and can be requested from the authors in the meantime.

## Supporting information

**Table S1:** T cell (T) and neutrophil (N) videos used in our project. Most videos captured cells moving in microchannels of varying width (2 µm to 30 µm). In four of the videos, cells moved in 6 µm microchannels with added 4-way splits and joins. We divided the videos into separate training and testing batches as shown.

| id | cells | geometry | frames | tracks | use |
| --- | --- | --- | --- | --- | --- |
| 1 | T | 6 $\mu\text{m}$ | 181 | 64 | training |
| 2 | T | 6 $\mu\text{m}$ | 181 | 70 | training |
| 3 | T | 6 $\mu\text{m}$ | 271 | 159 | training |
| 4 | T | 6 $\mu\text{m}$ | 271 | 124 | training |
| 5 | T | 6 $\mu\text{m}$ | 271 | 150 | training |
| 6 | T | 6 $\mu\text{m}$ | 180 | 185 | training |
| 7 | T | 6 $\mu\text{m}$ | 180 | 298 | training |
| 8 | T | 15 $\mu\text{m}$ | 271 | 111 | training |
| 9 | T | 15 $\mu\text{m}$ | 271 | 426 | training |
| 10 | T | 30 $\mu\text{m}$ | 271 | 753 | training |
| 11 | T | 30 $\mu\text{m}$ | 271 | 128 | training |
| 12 | T | 4-way | 361 | 56 | training |
| 13 | T | 4-way | 357 | 42 | training |
| 14 | N | 5 $\mu\text{m}$ | 271 | 64 | training |
| 15 | N | 5 $\mu\text{m}$ | 271 | 116 | training |
| 16 | N | 6 $\mu\text{m}$ | 271 | 122 | training |
| 17 | N | 6 $\mu\text{m}$ | 271 | 157 | training |
| 18 | T | 2 $\mu\text{m}$ | 271 | 267 | testing |
| 19 | T | 2 $\mu\text{m}$ | 181 | 34 | testing |
| 20 | T | 2 $\mu\text{m}$ | 271 | 28 | testing |
| 21 | T | 6 $\mu\text{m}$ | 180 | 379 | testing |
| 22 | T | 6 $\mu\text{m}$ | 180 | 254 | testing |
| 23 | T | 6 $\mu\text{m}$ | 181 | 86 | testing |
| 24 | T | 6 $\mu\text{m}$ | 271 | 115 | testing |
| 25 | T | 6 $\mu\text{m}$ | 271 | 214 | testing |
| 26 | T | 6 $\mu\text{m}$ | 180 | 551 | testing |
| 27 | T | 6 $\mu\text{m}$ | 180 | 139 | testing |
| 28 | T | 6 $\mu\text{m}$ | 271 | 30 | testing |
| 29 | T | 6 $\mu\text{m}$ | 271 | 72 | testing |
| 30 | T | 6 $\mu\text{m}$ | 180 | 152 | testing |
| 31 | T | 6 $\mu\text{m}$ | 271 | 39 | testing |
| 32 | T | 6 $\mu\text{m}$ | 271 | 195 | testing |
| 33 | T | 6 $\mu\text{m}$ | 180 | 455 | testing |
| 34 | T | 6 $\mu\text{m}$ | 180 | 90 | testing |
| 35 | T | 6 $\mu\text{m}$ | 180 | 82 | testing |
| 36 | T | 6 $\mu\text{m}$ | 180 | 181 | testing |
| 37 | T | 6 $\mu\text{m}$ | 180 | 127 | testing |
| 38 | T | 15 $\mu\text{m}$ | 271 | 111 | testing |
| 39 | T | 30 $\mu\text{m}$ | 271 | 742 | testing |
| 40 | T | 4-way | 541 | 21 | testing |
| 41 | T | 4-way | 541 | 28 | testing |
| 42 | N | 5 $\mu\text{m}$ | 271 | 31 | testing |
| 43 | N | 5 $\mu\text{m}$ | 271 | 104 | testing |
| 44 | N | 5 $\mu\text{m}$ | 271 | 15 | testing |
| 45 | N | 5 $\mu\text{m}$ | 271 | 191 | testing |
| 46 | N | 5 $\mu\text{m}$ | 271 | 670 | testing |
| 47 | N | 6 $\mu\text{m}$ | 180 | 181 | testing |
| 48 | N | 6 $\mu\text{m}$ | 271 | 45 | testing |
| 49 | N | 6 $\mu\text{m}$ | 180 | 210 | testing |
| 50 | N | 6 $\mu\text{m}$ | 271 | 139 | testing |
| 51 | N | 6 $\mu\text{m}$ | 180 | 207 | testing |
| 52 | N | 6 $\mu\text{m}$ | 180 | 155 | testing |
| 53 | N | 6 $\mu\text{m}$ | 180 | 228 | testing |
| 54 | N | 6 $\mu\text{m}$ | 120 | 122 | testing |
| 55 | N | 6 $\mu\text{m}$ | 180 | 102 | testing |
| 56 | N | 6 $\mu\text{m}$ | 180 | 231 | testing |

**Figure S1:**
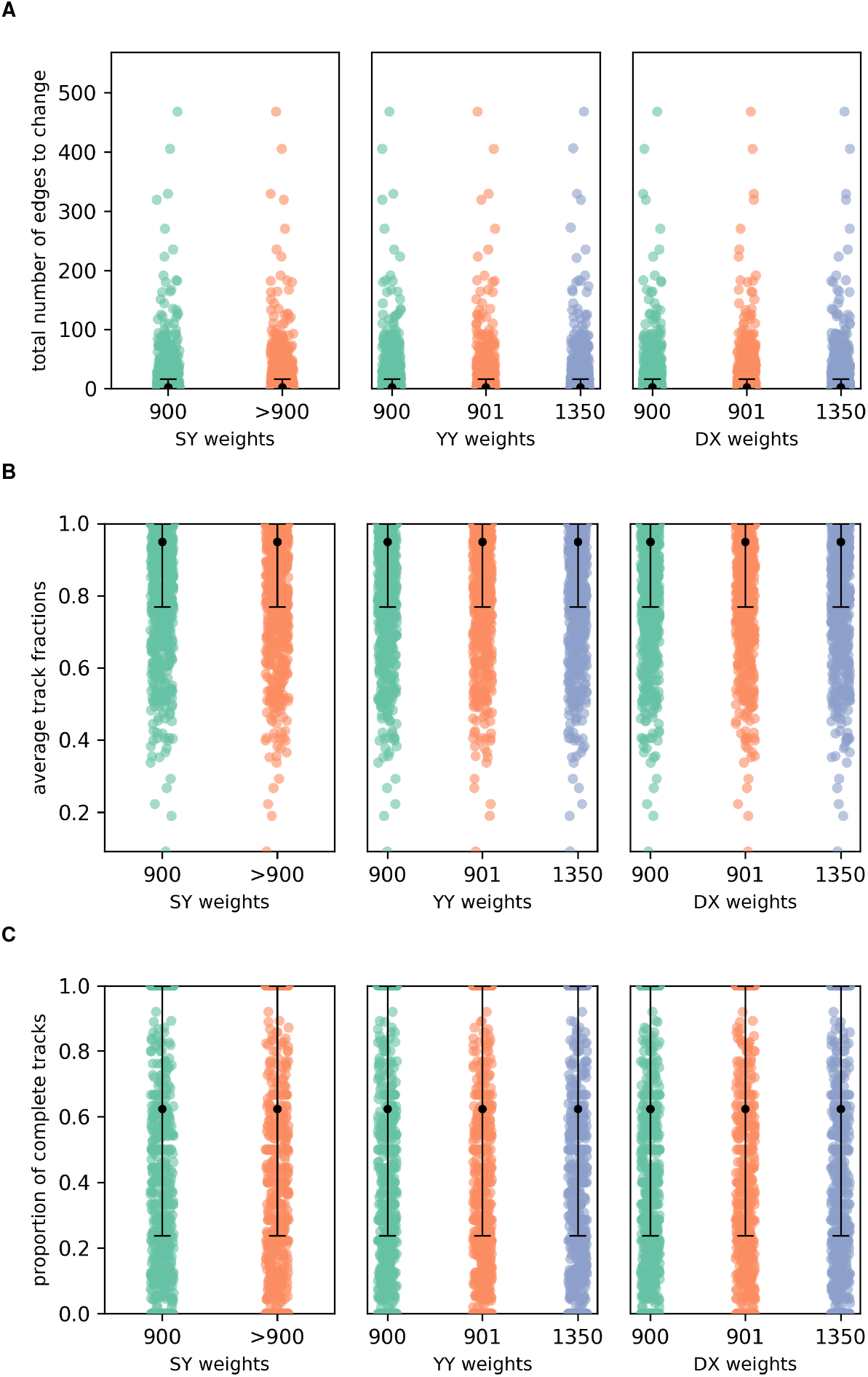
Experiments with edge weights for the edge types SY, YY and DX. The weights of the SY-, YY-, and DX-edges do not appear to affect the quality of the tracks for values greater than the maximum edge weight threshold, using a squared Euclidean distance without displacement prediction. **(A)** The total number of edges that need to be changed (edges to add + edges to remove) on a complete set of tracks generated by a single run of Dijkstra’s algorithm, compared to ground truth. **(B)** Average track fractions, calculated per channel. **(C)** Proportion of complete tracks, calculated per channel.

